# Hotmelt tissue adhesive with supramolecularly-controlled sol-gel transition for preventing postoperative abdominal adhesion

**DOI:** 10.1101/2021.10.26.464756

**Authors:** Akihiro Nishiguchi, Hiroaki Ichimaru, Shima Ito, Kazuhiro Nagasaka, Tetsushi Taguchi

**Affiliations:** Polymers and Biomaterials Field, Research Center for Functional Materials, National Institute for Materials Science, 1-1 Namiki, Tsukuba, Ibaraki 305-0044 (Japan); Faculty of Pure and Applied Sciences, University of Tsukuba, 1-1-1 Tennodai, Tsukuba, Ibaraki 305-8577 (Japan); Department of Chemistry, Faculty of Science Division I, Tokyo University of Science, 1-3 Kagurazaka, Shinjuku-ku, Tokyo 162-8601 (Japan)

**Keywords:** postoperative adhesion, tissue adhesive, supramolecular chemistry, gel, sol-gel transition

## Abstract

Postoperative adhesion is a serious and frequent complication, but there is currently no reliable anti-adhesive barrier available due to low tissue adhesiveness, undesirable chemical reactions, and poor operability. Here, we report a single-syringe hotmelt tissue adhesive to prevent postoperative abdominal adhesions. Through the augmentation of intermolecular hydrogen bonding by conjugation of the ureidopyrimidinone unit to tendon-derived gelatin, the sol-gel transition behavior of gelatin was supramolecularly-controlled, which provided a hotmelt tissue adhesive that dissolves upon warming over 40 °C and glues at 37 °C. This functionalization improved the key features necessary for an anti-adhesive barrier, including bulk mechanical strength, tissue adhesive properties, underwater stability, and anti-adhesive property. This hotmelt tissue adhesive with excellent tissue adhesiveness, biocompatibility, and operability has enormous potential to prevent postoperative complications.

## Introduction

Postoperative adhesion is a fibrous band of scar tissue between organs and tissues and is caused by injury after surgery. Its occurrence has been observed in more than 93% of the patients undergoing laparotomy.^[1]^ Despite recent progress in laparoscopic surgery, which is known to reduce the adhesion rate,^[2]^ postoperative adhesion remains a serious and frequent complication in clinical practice. Postoperative adhesions cause sequential severe complications, including bowel obstruction, infertility, and chronic pelvic pain^[3]^, which leads to poor postoperative quality of life, prolonged hospitalization, and readmission.^[4]^ In addition, postoperative adhesions increase the difficulty and time of the second surgery for adhesiolysis.

Postoperative adhesion is initiated by tissue injury caused by surgery, irritation, and trauma.^[5]^ The injury initiates secretion of vasoactive signaling molecules, such as histamines, resulting in an increase in the permeability of the blood vessels. Then, the coagulation cascade is stimulated to deposit fibrin and form clots. Infiltrating inflammatory cells secrete pro-inflammatory cytokines and promote extracellular matrix deposition.^[6]^ These excessive coagulation and inflammatory responses disrupt the balance between fibrin formation and fibrinolysis system, leading to the formation of adhesion between tissues and organs. To prevent postoperative adhesion, anti-adhesive barriers, such as film and liquid-type tissue adhesive, have been used clinically.^[7]^ The anti-adhesive barrier functions as a physical barrier under physiological conditions in order to protect injured tissues by adhering to and covering wet tissues and suppressing the penetration of fibrinogen and cells. Finally, the barrier degrades at some point and completes the wound healing process. A meta-analysis has shown that a film-type anti-adhesive barrier reduces the incidence of re-operation.^[8]^ However, the preventive effect is limited because it fails to tightly adhere to uneven irregular tissue surfaces and easily detaches after the swelling of the film.^[9]^ Moreover, a less flexible film lacks operability in laparoscopic surgery when delivered to injured tissues. In contrast, liquid-type tissue adhesives can be delivered using a double-syringe system and cover the tissues without depending on the geometry of the tissue surfaces.^[10]^ Most tissue adhesives are composed of two types of polymer solutions and form gels via chemical reactions such as the thiol-acrylate reaction^[11]^, amine-aldehyde reaction^[12]^, and amine-activated ester reaction^[13]^. Although chemical crosslinking can improve retention time of barriers on the tissues, chemically-crosslinked gels generally represent poor mechanical properties to follow the movement of tissues and non-specific chemical reactions may cause inflammatory responses. Moreover, the double-syringe system has risk of inefficient gelation due to the incomplete mixing of the two components. Despite their ability to prevent postoperative adhesion, the benefits and associated risks of these barriers remain unclear and these barriers have not been frequently used in abdominal surgery (<10%).^[8, 14]^

Recently, single syringe-type tissue adhesives have attracted increasing attention as highly biocompatible and operatable anti-adhesive barriers. A single-syringe tissue adhesive requires no chemical crosslinking reagents or mixing devices to apply adhesives to tissues.^[15]^ To date, polymer solutions (e.g., icodextrin) have been used to separate injured organs and tissues by instilling into the peritoneal cavity. However, polymer solutions currently show insufficient evidence for preventing postoperative adhesions.^[16]^ Non-gelling and non-adhesive polymer solutions may not be sufficient to separate injured tissues due to rapid clearance from the peritoneal cavity.^[17]^ To improve the tissue adhesive properties, several single-syringe adhesives, such as temperature-responsive polymer^[18]^, biomolecule-responsive adhesive^[19]^, and shear thinning gel system,^[20]^ have been studied. However, these adhesives have numerous associated concerns pertaining to tissue adhesiveness, mechanical properties, *in vivo* retention time under wet conditions, and therapeutic efficacy.

To develop a single-syringe tissue adhesive, we focused on a hotmelt adhesive. Hotmelt adhesive is a thermoplastic adhesive that is solid at room temperature, but becomes liquid when heated over the melting point, and then hardens to adhere to materials when returned to room temperature.^[21]^ The heat-induced phase transition of hotmelt adhesive enables an increase in the contact area with materials when heated and enhances the bulk mechanical strength of the adhesive when cooled, which contributes to an increase in both the interfacial strength and bulk strength. Leveraging the hotmelt system to tissue adhesives would provide a novel single-syringe adhesive for biomedical applications. However, in general, hotmelt adhesives are composed of non-biodegradable polymers, such as poly(ethylene vinyl acetate), which must be heated to 160°C or higher when used. Therefore, it is necessary to develop biodegradable tissue adhesives whose phase transition temperature is around body temperature, however, there have been no relevant reports published so far.

Here, we report the development of single-syringe, hotmelt-type, supramolecular gelatin-based tissue adhesives with controlled sol-gel transition behavior (Scheme 1). Gelatin, type of denatured collagen, possesses intrinsic sol-gel transition behavior in which the viscosity of the solution decreases above the transition temperature and gelation occurs below the temperature owing to the hydrogen bonding originating from collagen motifs. Although this sol-gel transition behavior is useful for engineering hotmelt tissue adhesives, commonly used porcine skin-derived gelatin (SG) exhibits a low transition temperature (~33 °C) and sol state at body temperature. We hypothesized that if the sol-gel transition temperature of gelatin is increased above body temperature (37 °C), the gelatin solution would be in sol state when warming up (>40 °C) and would form a gel at body temperature, which can be used as a hotmelt tissue adhesive. We selected porcine-derived tendon gelatin (TG), which has a higher sol-gel transition temperature than SG, and further functionalized TG with 2-ureido-4[1H]-pyrimidinone unit (UPy unit) to synthesize TG functionalized with UPy unit (TGUPy). UPy unit dimerizes through quadruple hydrogen bonding^[22]^ and can induce strong intermolecular non-covalent interactions between TGUPy, which may increase the sol-gel transition temperature of gelatin to achieve hotmelt tissue adhesives. We aimed to prevent postoperative abdominal adhesions using TGUPy-based adhesives (Scheme 1b). When TGUPy tissue adhesives were warmed up above the transition temperature (>40 °C) and applied to postoperative injury, liquid-like TGUPy could glue into small tissue gaps. TGUPy then transformed to gel at body temperature (37 °C), strongly adhering to the tissue, and covered injured surfaces. Once gelation was completed, the gel lost its adhesiveness toward other tissues and prevented abdominal adhesion during wound healing. This hotmelt tissue adhesive may serve as a medical material to prevent postoperative complications.

**Scheme 1.**
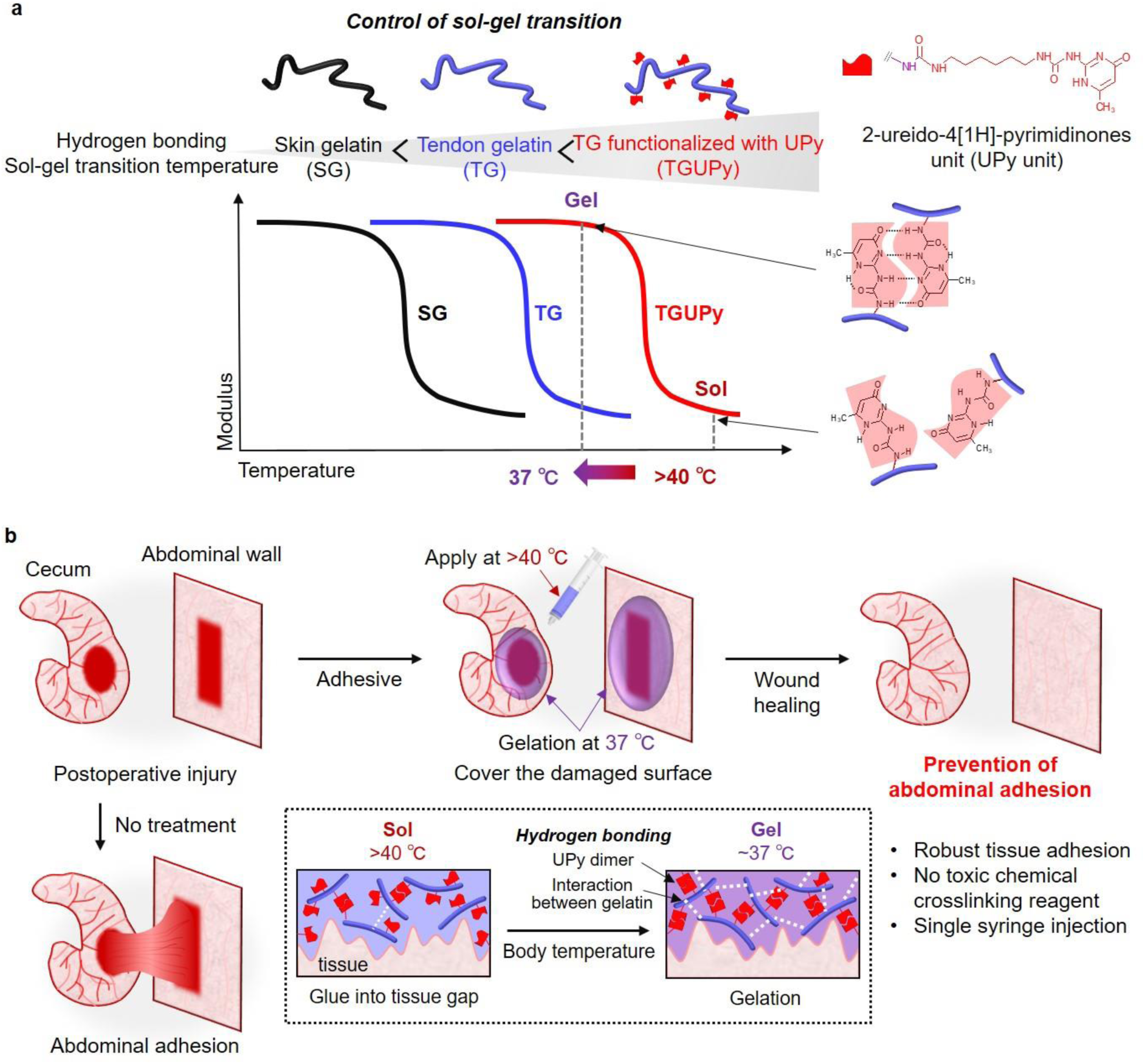
Schematic illustration of the design of hotmelt tissue adhesive and its operating principle. (a) Porcine-derived skin gelatin (SG) and tendon gelatin (TG) were used as a component of adhesive. TG was functionalized with 2-ureido-4[1H]-pyrimidinone unit (UPy unit) (TGUPy) to augment intermolecular hydrogen bonding. The degree of introduced hydrogen bonding and sol-gel transition temperature of gelatin were controlled based on supramolecular approach. By tuning sol-gel transition temperature, TGUPy hotmelt tissue adhesive, wherein the solution is in sol state above body temperature (37 °C) and becomes gel at 37 °C, can be obtained. (b) Prevention of abdominal adhesion using hotmelt tissue adhesive. TGUPy solution (sol state above 40 °C) was applied to postoperative injured tissues (e.g., cecum and abdominal wall). Low viscosity TGUPy above 40 °C can glue into tissue gaps and transform to gel at 37 °C through the formation of intermolecular hydrogen bonding. Gel covers injured surfaces and prevents postoperative adhesion during wound healing.

## Results and Discussion

### Temperature-dependent rheological properties of TGUPy gels

By controlling the intrinsic sol-gel transition behavior of gelatin through supramolecular approach, hotmelt tissue adhesives supramolecularly-crosslinked with complementary hydrogen bonding were developed. Gelatin exhibits reversible sol-gel transition behavior through hydrogen bonding originating from the collagen motif (glycine-X-Y, where X and Y are frequently proline (Pro) and hydroxyproline (Hyp)), and the sol-gel transition temperature depends on Pro and Hyp content.^[23]^ As shown in Table S1, the Pro and Hyp content in gelatin varies among species and body parts from where gelatin is derived. Among them, bovine and porcine-derived gelatin showed higher Pro and Hyp contents and melting points than fish-derived gelatin. In particular, TG showed the highest melting point (38.2 °C), partly because of the difference in collagen types and molecular weight.

Since UPy units are known to dimerize strongly through quadruple hydrogen bonding (dimerization constant *K*_dim_ > 10^6^ M^-1^ in CHCl_3_),^[22a]^ the conjugation of the UPy units may augment hydrogen bonding between gelatin and increase the sol-gel transition temperature. Moreover, the UPy unit dissociates when the temperature increases, which decreases the viscosity of the solution, similar to that observed for hydrogen bonding in gelatin. These functionalities based on hydrogen bonding would be favorable for the application of hotmelt tissue adhesives. The UPy unit was synthesized by the reaction of 2-amino-4-hydroxy-6-methylpyrimidine with 1,6-diisocyanatohexane (C6). TGUPy was synthesized through the formation of urea bonds between the TG and UPy units. The degree of substitution (D.S.) of the UPy unit was estimated from the measurement of residual amino groups because the reactivity of the primary amine was 1000 times faster than that of other functional groups such as carboxylic acid and hydroxyl groups.^[24]^ D.S. varied from 26% to 76% (mol% to amino groups in TG) (Table S2). TGUPy-26, -42, and -53 were soluble in phosphate buffered saline (PBS), whereas TGUPy with more than 79% D.S. was not soluble in PBS due to strong intra-/inter-molecular hydrogen bonding. The introduction of UPy onto TG was characterized by ^1^H-nuclear magnetic resonance (^1^H-NMR) spectroscopy (Figure S1). Fourier transform infrared (FT-IR) spectroscopy showed the disappearance of the peak at 2280 cm^-1^, attributed to isocyanate groups in TGUPy (Figure S2).

The solutions of TG and TGUPy with D.S. 42 (TGUPy-42) (20 wt% in PBS) were in the sol state at 45 °C (over the transition temperature). At 37 °C, TGUPy formed viscoelastic, adhesive gels, while TG was in a sol state with low viscosity (Figure 1a and Movie S1). The temperature-dependent rheological properties were evaluated using a rheometer to quantify the sol-gel transition behavior of TGUPy. The shear storage modulus (*G*’) of SG dropped above 30 °C and the sol-gel transition temperature was estimated to be 33.1 °C (Figure 1b). Of note, the sol-gel transition temperature was defined as the temperature at which the value of *G’* was lower than that of the shear loss modulus (*G*”). TG showed a higher transition temperature (38.2 °C) than SG and formed gels at 37 °C. TGUPy-42 showed the highest transition temperature (40.0 °C) and *G*’ (834 Pa) at 37 °C showed a 72-fold and 5.8-fold increase compared to that of SG and TG, respectively. Moreover, the sol-gel transition temperature and *G’* increased with increasing D.S. of the UPy unit in TG (Figure 1c). In contrast to TGUPy, SG functionalized with UPy unit (SGUPy) did not form gels at 37 °C (*G*’<*G*”), and the transition temperature did not exceed 37 °C even at high D.S. (Figure S3). These results suggest that functionalization of TG with UPy unit increased the sol-gel transition temperature by augmenting intermolecular hydrogen bonding, which may impart the hotmelt tissue adhesive property to gelatin.

**Figure 1.**
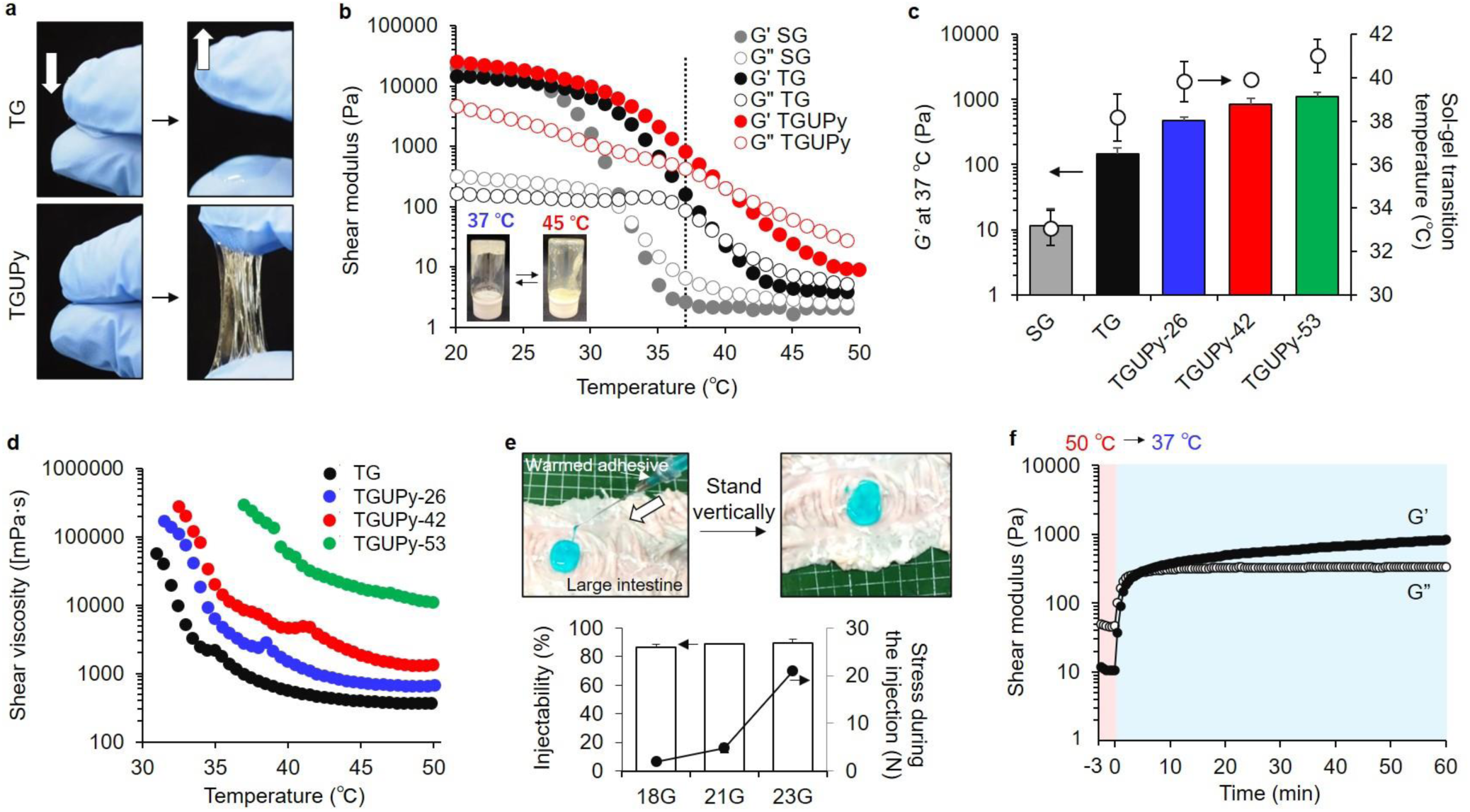
Rheological characteristics of TGUPy gels. (a) Macroscopic photos of TG and TGUPy. TG and TGUPy were warmed up at 45 °C and kept at 37 °C on fingers. TGUPy formed adhesive gels, while TG was at low viscosity and in sol state. (b) Temperature-dependent shear modulus of SG, TG, and TGUPy-42 solution dissolved in PBS at 20 wt%. The measurement of each sample was initiated at 20 °C with an increase in the temperature at the rate of 1 °C/min and the measurement was stopped when temperature reached 50 °C. (c) Effect of D.S. of UPy unit on *G’* and sol-gel transition temperature. The sol-gel transition temperature denotes the temperature when the value of *G’* was lower than that of *G”*. (d) Temperature-dependent shear viscosity of TG, TGUPy-26, -42, and -53. (e) Macroscopic photos of TGUPy-42 adhesive injected into a large intestine. The adhesive was injected with 18G-needle at 50 °C. After the incubation at 37 °C, gelation occurred and the gel adhered to vertically-standing tissue. (f) Gelation kinetics of TGUPy-42 gel when the temperature changed from 50 °C to 37 °C. Data are presented as mean ± s.d. (n = 3).

Although TGUPy-53 showed the highest sol-gel transition temperature and *G*’, TGUPy-53 was highly viscous even at 50 °C (>10000 mPa·s) (Figure 1d). Shear viscosity is an important parameter for the injection of adhesives, and TGUPy-53 was observed to be too viscous to be injected with a syringe. The shear viscosity of TGUPy-42 at 50 °C was 1350 mPa·s and TGUPy-42 was injectable withh a syringe. The injected adhesive rapidly formed gels at 37 °C and adhered to the porcine large intestine tissue (Figure 1e). TGUPy-42 was injected using 18, 21, and 23G-needle at high injectability and the required injection force was less than 21 N which is injectable range for an operator.^[25]^ In the following experiment, TGUPy-42 was used unless specified otherwise. The swelling ratio of the TGUPy-42 gel was 8.9 ± 0.3. The concentration of gelatin substantially affected *G*’ at 37 °C and 20 wt% TGUPy-42 showed the highest mechanical strength (Figure S4).

The gelation kinetics were determined by measuring the temperature-responsive rheological changes of TGUPy-42. When the temperature was changed from 50 °C to 37 °C, *G*’ quickly increased, and the gelation time was approximately 5 min (Figure 1f). *G*’ reached a plateau in hours. Even after the gelation, supramolecularly-crosslinked TGUPy adhesives exhibited reversible sol-gel transition behavior under cyclic temperature changes (Figure S5). The *G*’ at 50 °C decreased to less than one-tenth of that at 37 °C, and *G*’ increased to its original value when returned to 37 °C. The *G*’ value was maintained even after several cycles. Thus, this supramolecular approach to control the sol-gel transition behavior is useful for the design of hotmelt tissue adhesives.

### Structural properties of TGUPy gels

The mechanical strength of TGUPy gels was evaluated by tensile and compression tests. At 25 °C, TGUPy gels exhibited 1.7- and 4.3-fold increase in fracture strain (525%) and 2.8- and 10-fold increase in fracture strength (402 kPa) compared to those of SG and TG (Figure 2a). The same experiments were performed at 37 °C, but SG and TG were in the sol state at 37 °C and could not be transformed into gels. TGUPy formed viscoelastic gels at 37 °C and tensile test showed that the fracture strain and fracture strength were 294% and 200 kPa, respectively. Although the values at 37 °C were smaller than those at 25 °C, TGUPy maintained a stable structure at the body temperature. In the compression test, the TGUPy gels deformed under compression at 25 °C and maintained their original shapes against a maximum of 50 N of compression (Figure 2b,c). The SG and TG gels were brittle and broke after the compression test. At 37 °C, the TGUPy gels were softened and deformable, but not brittle. These results suggest that the functionalization of TG with the UPy unit substantially improved the mechanical properties of the gels. It has been reported that the UPy unit dimerized by quadruple hydrogen bonding can dissipate external energy under a large deformation.^[26]^ In TGUPy gels, hydrogen bonding of the UPy unit may work as a sacrificial bond and dissipate external energy. Moreover, the reversible formation of hydrogen bonding in TGUPy gels possibly rearranged the configuration of the polymer network under deformation, which prevented crack propagation and in turn toughened the gels. Moreover, rheological analysis of TGUPy gels at 37 °C revealed that the gels exhibited solid-like behavior with linear viscoelastic responses in strain-dependent rheological measurements (Figure S6a). Because the highest strain applied to the tissues in the body is estimated to be approximately 10%,^[27]^ the gels are expected to maintain a stable solid-like structure under deformation. Moreover, TGUPy gels showed frequency response observed in supramolecular gels and solid-like structures in the range of 1 to 100 rad s^-1^ (Figure S6b). These results indicate that TGUPy adhesives with non-covalent bonding networks possessed viscoelastic properties under physiological conditions.

**Figure 2.**
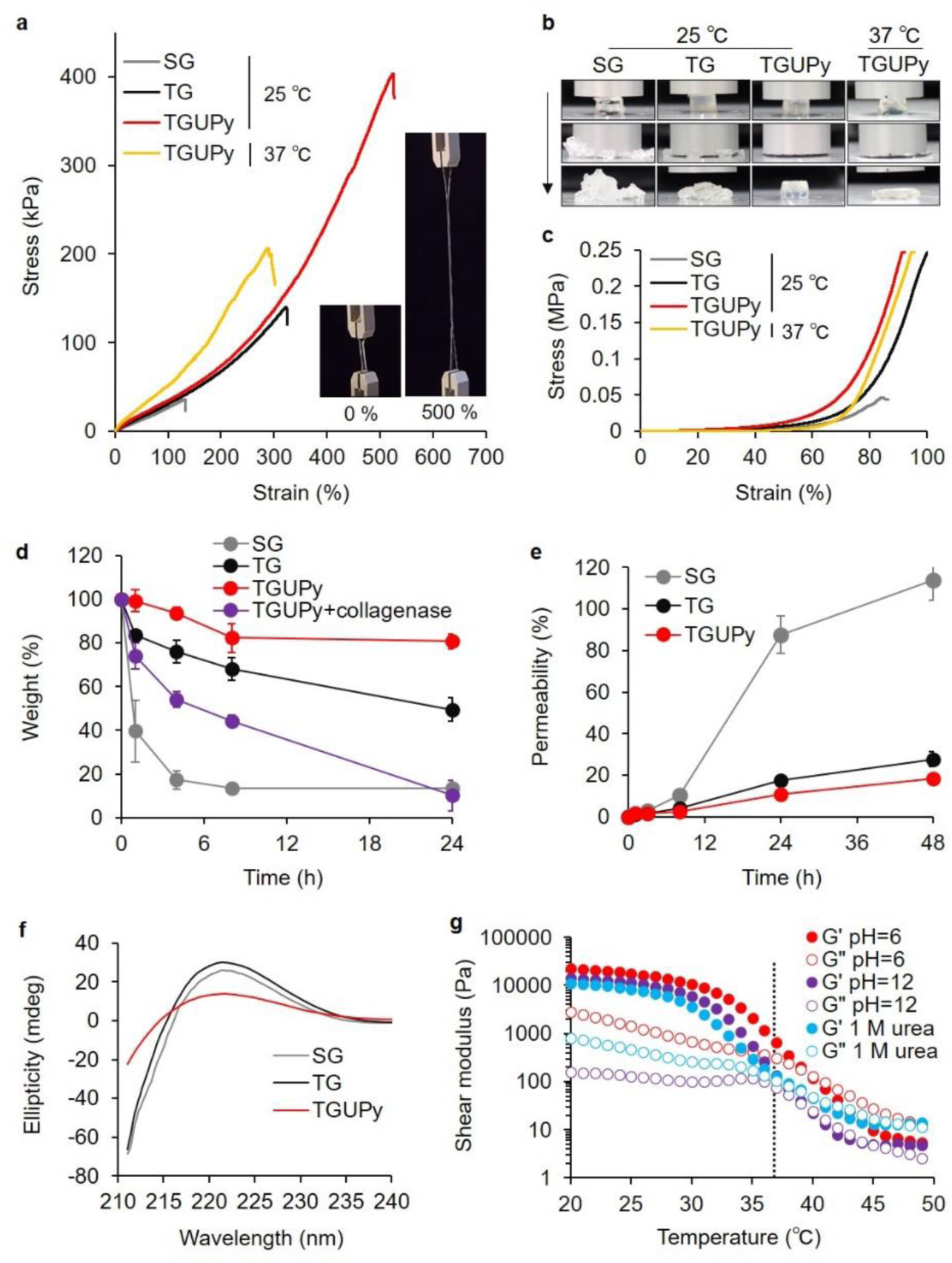
Structural properties of TGUPy gels. (a) Tensile test of SG, TG, and TGUPy-42 gels (20 wt%, PBS). The measurement was performed at 25 and 37 °C. The insets showed images of TGUPy-42 gels during tensile test. (b,c) Macroscopic images and compression test of SG, TG, and TGUPy-42 gels. The measurement was performed at 25 and 37 °C. (d) Underwater stability test of SG, TG, and TGUPy-42 gels with and without collagenase. Each gel was incubated in PBS with and without collagenase (120 U/mL) at 37 °C. (e) Dye transfer test of SG, TG, and TGUPy-42 gels. Fluorescently-labeled albumin (0.1 mg/mL) was used as a model protein. (f) CD spectroscopic measurement of SG, TG, and TGUPy-42 gels. (g) Temperature-dependent shear modulus change of TGUPy-42 gels (pH=6 and 12 in PBS) and TGUPy-42 gels with 1M urea. Data are presented as mean ± s.d. (n = 3).

Underwater stability is a crucial characteristic in the application of tissue adhesives that work under wet conditions. When SG gels were exposed to PBS at 37 °C, SG completely degraded after 24 h because of its low transition temperature of ~33 °C (Figure 2d). TGUPy gels exhibited higher underwater stability compared to TG gels (81% vs. 50% at 24 h), indicating that supramolecular crosslinking with UPy units suppressed the dissolution of gelatin and improved underwater stability. In contrast, based TGUPy gels were enzymatically degraded in the presence of collagenase, indicating that the TGUPy gels were biodegradable. Moreover, the barrier function of gels as an anti-adhesive barrier was assessed by monitoring the transfer of fluorescently labeled albumin (*M*_w_ = 66.5 kDa), which was used as a model protein (Figure 2e). Postoperative adhesion is promoted by the leakage of fibrinogen to form a fibrin scaffold, and hence, the anti-adhesive barrier needs to possess barrier function against proteins. The SG gels dissolved during the incubation and showed high permeability. The transfer of albumin across the gels was greatly suppressed in TGUPy gels compared to TG gels. TGUPy gels possibly formed densely crosslinked polymer networks, which may prevent the leakage of fibrinogen from injured tissues. These results suggest that UPy-functionalized gelatin provides not only a hotmelt property but also improved mechanical strength, underwater stability, and barrier function, which would be favorable for its application as an anti-adhesive barrier.

Circular dichroism (CD) spectroscopic measurements were performed to understand the gelation mechanism of TGUPy (Figure 2f). CD spectroscopy is sensitive to the secondary structure of proteins and is useful for determining the helical structure of gelatin originating from collagen triple helices.^[28]^ CD measurements revealed that the peak at 221 nm ascribed to collagen triple helix formation between gelatin decreased in TGUPy solution compared to that in SG and TG solution. This result indicates that UPy functionalization attenuates the interaction between the helices of gelatin, partly due to bulky UPy units. Given that G’ at 37 °C and sol-gel transition temperature were increased by UPy-functionalization, the contribution of hydrogen bonding of UPy units may be large enough to improve the mechanical properties. To confirm whether the hydrogen bonding of UPy works as a crosslinking point, a blocking test using NaOH and urea was performed (Figure 2g). Upon increasing the pH to 12 or upon addition of urea, G’ of TGUPy gels decreased to almost the same value as that of the TG gels (Figure S7). The protonation of the UPy unit upon an increase in the pH and competitive inhibition of the dimerization of the UPy unit upon addition of urea suppressed the formation of hydrogen bonds between UPy units. This result indicates that hydrogen bonding is the main driving force for improving the structural properties of the TGUPy adhesives.

### Tissue adhesive properties

To apply a hotmelt tissue adhesive to an anti-adhesive barrier, the control of tissue adhesive property is pivotal, where it should adhere and should not. We aimed to develop hotmelt tissue adhesives that would (1) adhere to injured tissue surfaces when injected and (2) lose adhesiveness toward other tissues after gelation. To address the tissue adhesiveness of TGUPy adhesives, an *ex vivo* adhesion test was performed using porcine large intestine according to the American Society of Testing and Materials (ASTM) procedure (ASTM F-2258-05, Standard Test Method for Strength Properties of Tissue Adhesives in Tension) (Figure 3a). Adhesives pre-warmed at 50 °C were applied to the serosa of the large intestine (37 °C) and immediately pressed with another tissue (adhesion test). Note that the temperature of the adhesive rapidly dropped from 50 °C to 38 °C within 30 s when placed on the tissues, suggesting that this procedure may not damage the tissues (Figure S8 and Movie S2). When pulling apart the tissues, TGUPy was observed to have strongly adhered to the tissues and stretched gel was observed, while SG dissolved during the incubation and TG did not show strong tissue adhesion (Figure 3b and Movie S3). Adhesion test revealed that TGUPy-26 and -42 gels possessed higher tissue adhesion strength than those of SG and TG (Figure 3c,d). Because TGUPy-26 and -42 showed high *G’* at 37 °C as well as higher sol-gel transition temperature as shown in Figure 1c, improved bulk mechanical properties may enhance tissue adhesiveness. The fluidity of TGUPy-26 and -42 at 50 °C was sufficient to glue into tissue gaps, leading to a large contact area between the tissue and adhesive and high interfacial adhesion strength. SG was in the sol state at 37 °C and did not work as an adhesive. Although TG possessed sufficient fluidity to glue into the tissue gaps, the bulk mechanical strength was low. TGUPy-53 showed the highest *G’* and sol-gel transition temperatures, but the viscosity at 50 °C was too high to glue into the tissue gaps. Histological images of hematoxylin-eosin (HE)-stained tissues after the adhesion test showed that cohesive fracture occurred in TG, TGUPy-26, and -42 gels, indicating that each gel strongly glue the top and bottom tissues with high interfacial adhesion (Figure 3e and Figure S9). On the other hand, fluidic SG and highly viscous TGUPy-53 did not adhere to the top tissues. These results have highlighted the importance of both the bulk mechanical strength of gels at body temperature for cohesion force and fluidity before gelation for interfacial adhesion to develop hotmelt tissue adhesives.

**Figure 3.**
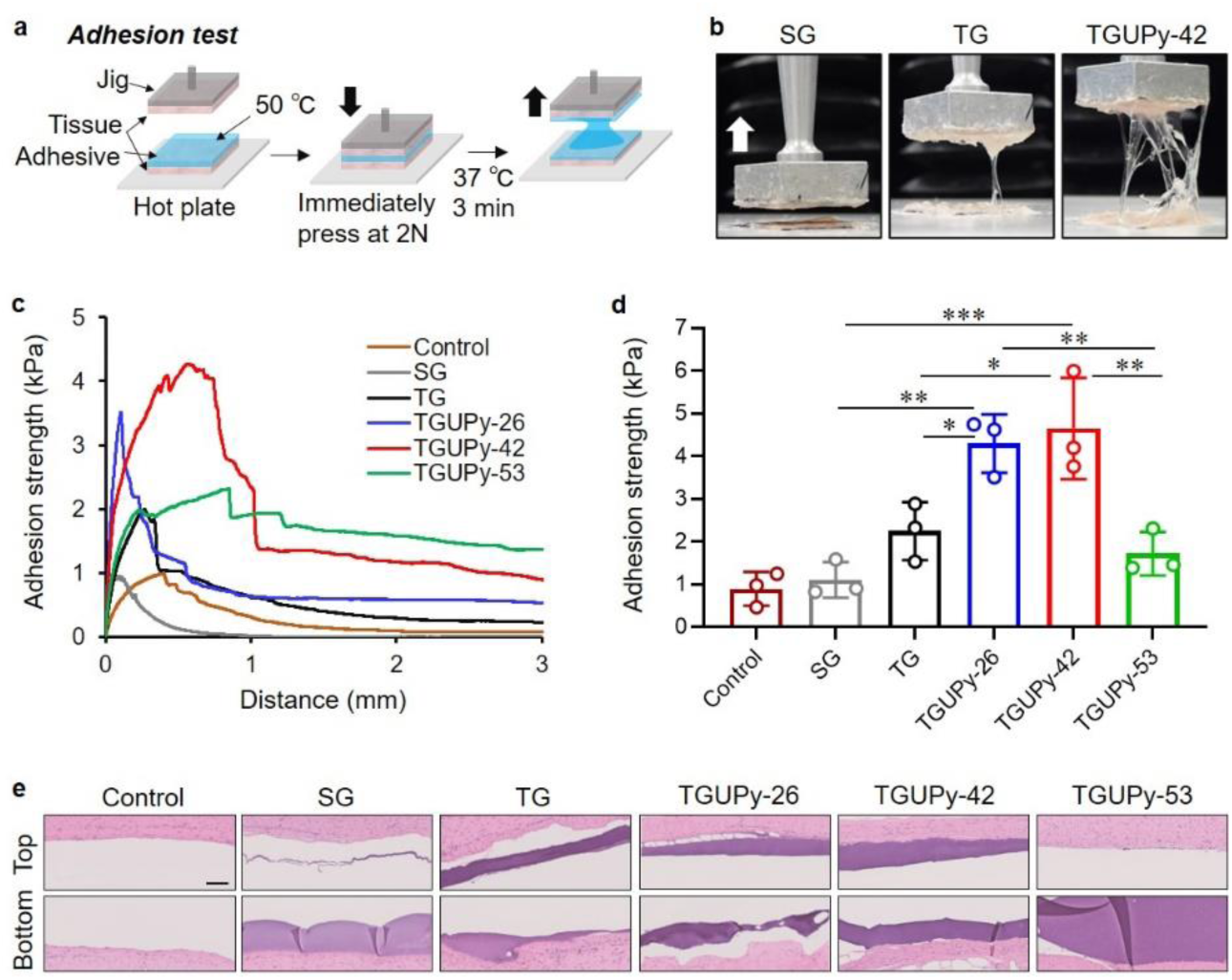
Ex vivo adhesion test. (a) Adhesion strength measurement using porcine large intestine according to ASTM F-2258-05. Adhesive at 50 °C was applied to the tissues and immediately pressed at 2 N for 3 min at 37 °C and the tensile strength was measured. (b) Macroscopic images of tissues with SG, TG, and TGUPy-42 during the test. (c,d) Adhesion strength of SG, TG, TGUPy-26, -42, and -53 to the tissues. Control denotes the adhesion test of tissues without gels. (e) HE images of the top and bottom tissues with SG, TG, TGUPy-26, -42, and -53 after the adhesion test. Data are presented as mean ± s.d. (n = 3). \**P* < 0.05, \*\**P* < 0.01, and \*\*\**P* < 0.001, analyzed by one-way analysis of variance (ANOVA) with Tukey’s multiple comparison test. Scale bar represents 100 μm.

Furthermore, we examined the effect of spacer molecules in the UPy unit on tissue adhesion. In addition to C6, 1,4-diisocyanatobutane (C4) and 1,8-diisocyanatoctane (C8) were used to synthesize UPy units with C4 and C8 spacers (D.S. = 42–46). The introduction of C4- and C8-UPy units into the TG was characterized by ^1^H NMR (Figure S10). By increasing the length of spacer molecules in TGUPy, *G’* at 37 °C as well as the shear viscosity increased (Figure S11a,b). Tissue adhesiveness was almost the same among C4-, C6-, and C8-TGUPy (Figure 11c). These results indicate that spacer length influences the rheological properties and mobility of UPy units, but does not change the adhesive strength.

### Anti-adhesive properties

The tissue adhesive itself should not cause abdominal adhesions. If the gelation process is incomplete, the adhesive may adhere to other undesirable tissues. To check whether TGUPy gels cause adhesion, *ex vivo* anti-adhesion tests were performed in the same manner as the adhesion test, except for the difference in the temperature of the adhesive when pressed. The adhesives warmed up to 50 °C were applied to the tissues and incubated at 37 °C for 10 min to complete gelation (Figure 4a). Two pieces of tissue were pressed, and adhesion strength was measured. The results of the anti-adhesion tests revealed that the top tissues did not adhere to any TGUPy gel at 37 °C (Figure 4b). TG partially adhered to the top tissues and showed the highest adhesion strength (Figure 4c). All TGUPy gels showed low adhesion strength at 37 °C because the sol-gel transition temperature was much higher than 37 °C and the gelation was complete. We evaluated the ratio of adhesion strength at 50 °C / adhesion strength at 37 °C to compare its ability as an anti-adhesive barrier. TGUPy-26 and -42 showed higher scores than TG and TGUPy-53 (Figure 4d). Moreover, HE images showed that TGUPy-42 and TGUPy-53 did not adhere to the top tissues, while TG and TGUPy-26 partly adhered to the top tissues (Figure 4e and Figure S12). UPy-functionalization enhanced the sol-gel transition temperature and structural integrity of gels through hydrogen bonding during UPy dimerization, leading to anti-adhesiveness. These results suggest that the TGUPy-42 adhesive, with robust tissue adhesiveness and excellent anti-adhesive properties, can be used to prevent postoperative adhesion.

**Figure 4.**
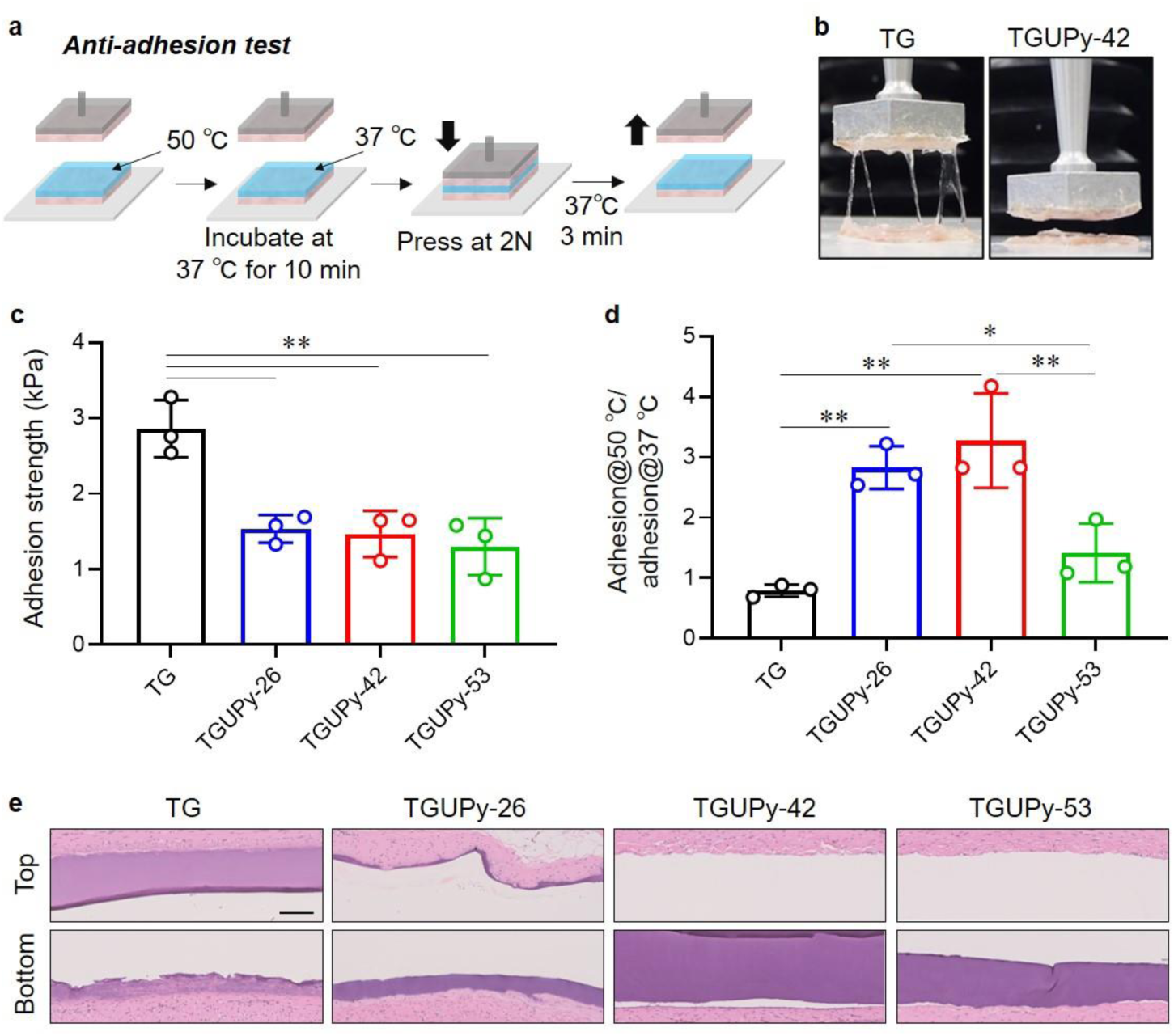
Ex vivo anti-adhesion test. (a) Anti-adhesion strength measurement using porcine large intestine according to ASTM F-2258-05. Adhesive at 50 °C was applied to the tissues and incubated for 3 min at 37 °C. The top tissue was then pressed at 2 N for 3 min at 37 °C and tensile strength was measured. (b) Macroscopic images of tissues with TG and TGUPy-42 during the test. (c) Adhesion strength of TG, TGUPy-26, -42, and -53 to the tissues. (d) Ratio of adhesion strength at 50 °C and adhesion strength at 37 °C of TG, TGUPy-26, -42, and -53. (e) HE images of the tp and bottom tissues with TG, TGUPy-26, -42, and -53 after anti-adhesion test. Data are presented as mean ± s.d. (n = 3). \**P* < 0.05 and \*\**P* < 0.01, analyzed by one-way ANOVA with Tukey’s multiple comparison test. Scale bar represents 100 μm.

### Prevention of abdominal adhesion

To confirm the biocompatibility of TGUPy adhesives, a cytocompatibility test was performed using cultured L929 mouse fibroblasts prior to *in vivo* experiments. We confirmed the high biocompatibility of TG and TGUPy in cytotoxicity assay (Figure S13). Moreover, each adhesive was subcutaneously implanted into mice to assess the biocompatibility and biodegradability. HE images revealed that SG and TG were resorbed at day 1 after the implantation, while TGUPy gels maintained their structure on day 1 owing to the high underwater stability of gels supramolecularly-crosslinked with hydrogen bonding of the UPy units (Figure S14). At day 3, all adhesives were resorbed, and there were few inflammatory cells on day 7. It has been reported that fibrinous exudates secreted after the injury are transient and broken by fibrinolysis within 72 h.^[29]^ Therefore, 72 h post-injury is a crucial time point that decides whether the abdominal adhesion occurs or not. TGUPy gels maintained their structure after 24 h *in vivo*, and *in vivo* retention might be prolonged by UPy-functionalization. TGUPy gels were degraded within 7 days and may not interfere with the wound healing process. Therefore, we expected that the degradation rate of TGUPy gel would be suitable for application as an anti-adhesive barrier.

Finally, we addressed the anti-adhesive ability of hotmelt tissue adhesives using cecum-abdominal wall adhesion models of rats (Figure 5a). To prepare the abdominal adhesion models, the cecum was scratched using gauze, and the surface of the abdominal wall was dissected using a scalpel. TG and TGUPy adhesives warmed up to 45 °C were applied to the injured cecum and abdominal wall, covering the tissues (Figure 5b). As a clinically-used anti-adhesive barrier, Seprafilm (Genzyme), which is a transparent film composed of hyaluronic acid and carboxymethyl cellulose, was used. TGUPy transformed from sol to gel on the tissue surfaces at 37 °C within a few minutes and strongly adhered to the tissues. At day 14 after the injury, untreated cecum and abdominal wall (control) caused severe adhesion, and the adhesion score was 3.5 (Figure 5c). While moderate abdominal adhesions were observed in TG-treated models, rats treated with the TGUPy adhesive and Seprafilm exhibited no abdominal adhesions (Figure 5d). The TGUPy adhesive exhibited a significant difference as compared to TG (score: 0 vs. 1.63). This result indicates that the TGUPy adhesive possessed anti-adhesive ability and was comparable to the clinically used anti-adhesive barrier.

**Figure 5.**
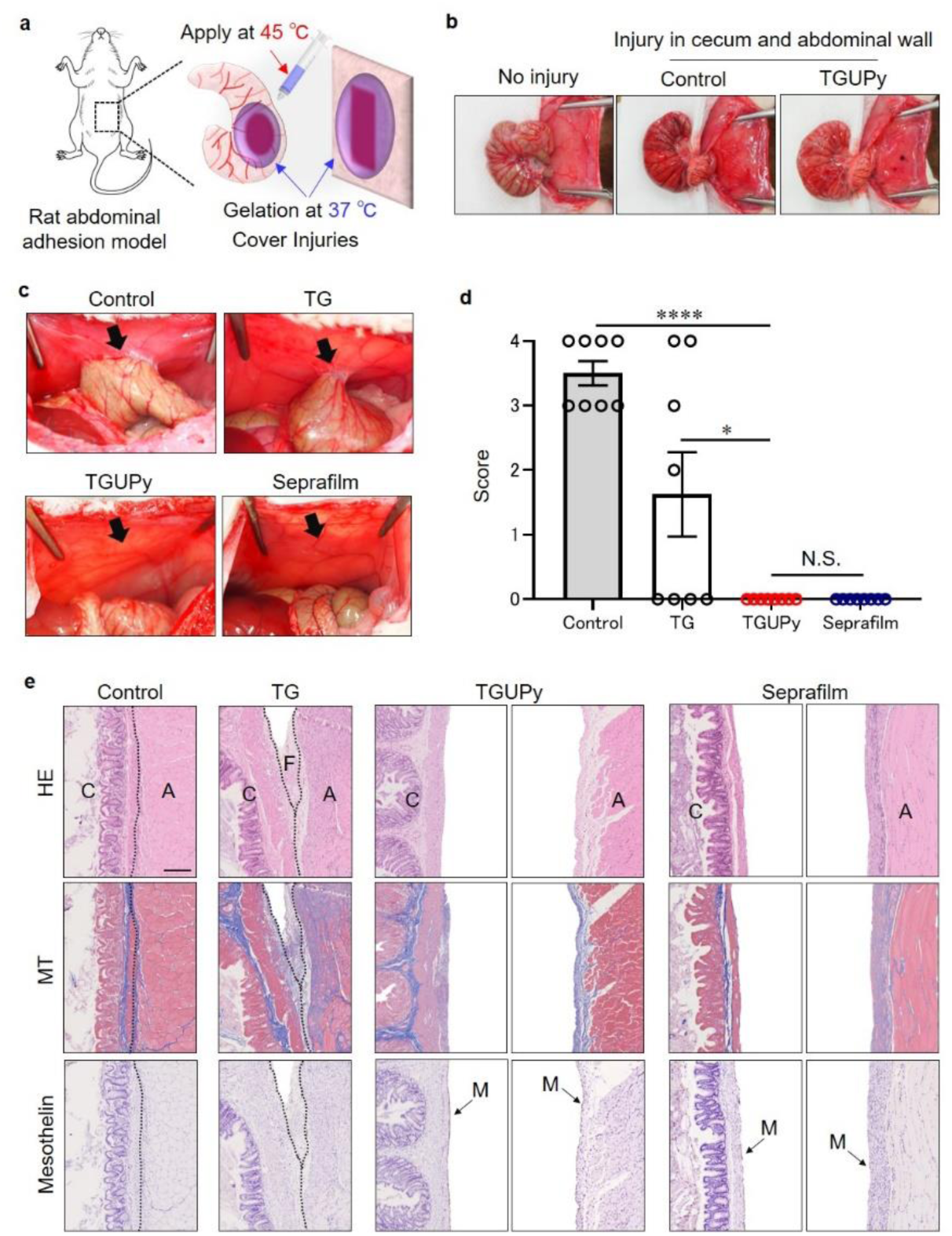
In vivo anti-adhesion test using cecum-abdominal wall adhesion models of rats. (a) Cecum and abdominal wall of rats were injured and treated with TG and TGUPy adhesives and Seprafilm. Adhesives were warmed up to 45 °C (sol state) and applied to injured tissues; they formed gel layer at 37 °C (gel state). (b) Macroscopic photos of cecum and abdominal wall prior to the injury (no injury) and after the injury with bleeding and treated with TG and TGUPy-42 adhesives and Seprafilm. Control denotes no treatment. (c) Macroscopic photos of cecum-abdominal wall adhesion in control, TG, TGUPy, and Seprafilm-treated samples. (d) Score of abdominal adhesion. The severity of adhesion was classified as follows: score 0: no adhesions, score 1: mild adhesions, score 2: localized moderate adhesions, score 3: moderate and wide adhesions, and score 4: severe adhesions, impossible to separate. (e) Histological observation of cecum and abdominal wall tissues stained with HE, Mason trichrome (MT), and mesothelin antibody. C: cecum, A: abdominal wall, F: fibrous tissue, M: mesothelial layer. Data are presented as mean ± s.e.m. (n = 8). \**P* < 0.05, \*\*\*\**P* < 0.0001, analyzed by one-way ANOVA with Tukey’s multiple comparison test. Scale bar represents 200 μm.

Histological observation showed that in control models, severe adhesion between the cecum and abdominal wall occurred and a thin fibrous tissue layer was formed, while the TG-treated cecum and abdominal wall partially adhered and formed fibrous tissues wherein fibrin network, fibroblasts, and immune cells were observed (Figure 5e). Sol-like TG may lack tissue adhesiveness and underwater stability under physiological conditions (37 °C, wet) and did not prevent abdominal adhesions. TGUPy-treated models showed no adhesion between the cecum and abdominal wall, and severe fibrosis was not observed with Masson’s trichrome (MT) staining. TGUPy adhesives were not observed on the tissues 14 days after the implantation, indicating the degradation of gels. Moreover, the mesothelial layer, which covers the organ surfaces and is responsible for separating organs, was recovered on the cecum and abdominal wall in TGUPy-treated rats. These results indicate that TGUPy adhesives protect injured tissue surfaces, suppress inflammatory responses, and do not interfere with the migration of mesothelial cells. This single-syringe hotmelt tissue adhesive can be considered as a powerful medical intervention to prevent postoperative adhesion and promote tissue regeneration.

Leveraging hotmelt-based adhesion through supramolecular approach enabled to engineer medical tissue adhesives with excellent tissue adhesiveness, biocompatibility, and operability. Robust tissue adhesion was achieved through the unique adhesion mechanism, wherein the fluidic adhesive glued the tissue gaps when warmed up and solidified with high mechanical properties when cooled to the body temperature. In terms of biocompatibility, there is no need to use toxic chemical crosslinking reagents for gelation. Gelatin is less immunogenic and has a low risk of infection and pyrogen contamination. Moreover, gelatin is derived from ECM proteins and expresses biofunctionality, including cell adhesion and biodegradability, which may serve as a scaffold to support wound healing. In terms of operability, pre-filled single-syringe hotmelt tissue adhesive is highly operable and can be immediately used in an emergency by warming up. Since two syringes for mixing two polymer components are not required, there is a low risk of insufficient mixing and clogging of the adhesive. We expect that the single-syringe adhesive would be more easily delivered to the injured tissues in laparoscopic surgery compared to anti-adhesive films. Moreover, liquid-based adhesives are independent of the roughness and size of injured tissues for adhesion and can be used to prevent postoperative adhesion in various organs and tissues, including the pelvic, uterine, intestinal, dural sac, tendon, and pericardium.^[30]^ Postoperative treatment with adhesives may reduce the risk of adhesion, pain, paralysis, and the cost of treatment and re-surgery.

Although many drugs, including anti-inflammatory drugs, anticoagulants, and fibrinolytics, have been studied to prevent postoperative adhesion, few have proven to be effective in clinical trials due to their rapid clearance from the peritoneal cavity and limited therapeutic efficacy.^[31]^ The combination of anti-inflammatory drugs with adhesives enables local delivery of drugs in a controlled-release manner, which may further improve the therapeutic efficacy. In addition to the anti-adhesive barrier, hotmelt tissue adhesive is available as a sealant, hemostatic reagent, and wound dressing, which can prevent various postoperative complications, including delayed bleeding, perforation, and inflammation.

For clinical application, further studies are required to optimize the TGUPy adhesive for higher therapeutic efficacy using a large animal model. Large-scale production with good manufacturing practices and detailed analysis of biodegradability, toxicology, and clearance of by-products need to be studied for future clinical translation.

### Conclusion

In summary, we have developed a hotmelt tissue adhesive with excellent tissue adhesiveness, biocompatibility, and operability to prevent postoperative abdominal adhesions. Supramolecular control of the sol-gel transition behavior of gelatin provided a hotmelt property in which fluidic sol over 40 °C glued into tissue gaps and formed gels at 37 °C, resulting in robust tissue adhesion through augmented hydrogen bonding between UPy units. UPy-functionalization increased the sol-gel transition temperature, bulk mechanical strength, and underwater stability under physiological conditions. Fluidic sol over the transition temperature was injected using a single syringe and rapidly formed gels when the temperature reached 37 °C. The TGUPy adhesive revealed high tissue adhesiveness toward the target tissues and excellent anti-adhesive property as compared to that of TG. Supramolecularly-crosslinked adhesives with UPy units do not require toxic chemical crosslinking reagents for gelation. *In vivo* experiments using cecum and abdominal wall adhesion models of rats showed that TGUPy adhesives prevented the abdominal adhesion. This approach overcomes the limitations of conventional anti-adhesive barriers and TGUPy adhesives may serve as a biomedical material that prevents postoperative complications and contributes to minimally invasive surgery.

## Experimental Procedures

### 1. Synthesis of UPy unit

2-ureido-4[1H]-pyrimidinone unit (UPy unit) was synthesized according to previous report.[1] Briefly, 2-amino-4-hydroxy-6-methylpyrimidine (33.0 mmol; 4.1 g, Sigma-Aldrich, USA) was dispersed in 1,6-diisocyanatohexane (148.6 mmol; 25.0 g, C6, Tokyo chemical Industry Co., Ltd., Japan) The reaction was initiated to heat the solution at 100 °C and continued for 16 h under stirring. After cooling to 25 °C, 10-volumes hexane was added to precipitate the product. The resulting precipitate was collected by filtration and washed with n-hexane three times. The product was dried at 50 °C under reduced pressure to obtain UPy units as a white powder. Yield: 90% (8.7 g, 29.7 mmol). UPy unit was characterized by ^1^H-nuclear magnetic resonance (^1^H-NMR, DMSO-d6, ECZ 400S, 400 MHz, JEOL, Japan). δ =11.53 (s, 1H, C*H*_3_–C–N<u>H</u>), 9.67 (s, 1H, C*H*_2_– NH–(C=O)–N<u>H</u>), 7.30 (t, 1H, C*H*_2_–N<u>H</u>–(C=O)–NH), 5.77 (s, 1H, C<u>H</u>=C–C*H*_3_), 3.12 (m, 4H, NH–(C=O)–NH– C<u>H</u>_2_ +C<u>H</u>_2_–NCO), 2.10 (s, 3H, C<u>H</u>_3_), 1.54 (m, 2H, NH–C*H*_2_–C<u>H</u>_2_), 1.45 (m, 2H, C<u>H</u>_2_–C*H*_2_– NCO), 1.31 (m, 4H, C*H*_2_–C*H*_2_–C<u>H</u>_2_–C<u>H</u>_2_–C*H*_2_–C*H*_2_– NCO) ppm.

For the synthesis of UPy units with different spacer length, 1,4-diisocyanatobutane (C4, Sigma-Aldrich, USA) and 1,8-diisocyanatoctane (C8, Sigma-Aldrich, USA) were used instead of C6. The synthesis procedure was the same manner as C6. UPy-C4: δ =11.52 (s, 1H, C*H*_3_–C–N<u>H</u>), 9.68 (s, 1H, C*H*_2_–NH–(C=O)–N<u>H</u>), 7.35 (t, 1H, C*H*_2_–N<u>H</u>–(C=O)–NH), 5.77 (s, 1H, C<u>H</u>=C–C*H*_3_), 3.16 (m, 4H, NH–(C=O)–NH–C<u>H</u>_2_ +C<u>H</u>_2_–NCO), 2.11 (s, 3H, C<u>H</u>_3_), 1.53 (m, 4H, NH–C*H*_2_–C<u>H</u>_2_) ppm. UPy-C8: δ =11.54 (s, 1H, C*H*_3_–C–N<u>H</u>), 9.64 (s, 1H, C*H*_2_–NH– (C=O)–N<u>H</u>), 7.39 (t, 1H, C*H*_2_–N<u>H</u>–(C=O)–NH), 5.77 (s, 1H, C<u>H</u>=C–C*H*_3_), 3.12 (m, 4H, NH–(C=O)–NH–C<u>H</u>_2_ +C<u>H</u>_2_–NCO), 2.10 (s, 3H, C<u>H</u>_3_), 1.54 (m, 2H, NH–C*H*_2_–C<u>H</u>_2_), 1.45 (m, 2H, C<u>H</u>_2_–C*H*_2_– NCO), 1.31 (m, 8H, C*H*_2_–C*H*_2_–C<u>H</u>_2_–C<u>H</u>_2_– C<u>H</u>_2_–C<u>H</u>_2_–C*H*_2_–C*H*_2_– NCO) ppm.

### 2. Synthesis of TGUPy

TGUPy was synthesized according to previous report with slight modification.[2] The 1g of porcine tendon-derived gelatin (TG, *M*_w_ = 344,000 Da, amino group: 293 μmol/g, Nitta Gelatin Inc., Japan) was dissolved in 20 mL of DMSO at 50°C under stirring. After cooling the solution to 25 °C, UPy unit (38.6 mg, 0.132 mmol, 45 mol% equivalent to amino groups in TG) was dispersed in 5 mL of DMSO and added to TG solution. The reaction was continued for 24 h at 25 °C under stirring. For the purification, the solution obtained was slowly added to 20-volumes of cold mixture solvent of ethanol and ethyl acetate (*v*/*v*=1/1) under stirring. The precipitates were then collected with a glass filter and washed with cold chloroform to remove unreacted UPy. After collecting the precipitates with the filtration, the samples were further washed with ethanol twice. The products were dried at room temperature under reduced pressure for 3 days. To synthesize TGUPy derivatives, different D.S. (30, 60, 90, and 120 mol% UPy) and spacer length (C4, C8) were used. The modification ratio of UPy in TG calculated by determining the residual amino groups using 2,4,6-trinitrobenzenesulfonic acid sodium salt dihydrate (TNBS, Tokyo chemical Industry Co., Ltd., Japan).[3] Briefly, 0.1 ml of 0.1% TG or TGUPy, 0.1 ml of 0.1% triethyl amine (Wako, Japan), and 0.1 ml of 0.1% TNBS were added to a 48-well plate and incubated at 37 °C for 2h. After adding 0.05 ml of 6 N HCl, the absorbance at 340 nm was measured using a microplate reader (Spark10M, TECAN, Switzerland). Calibration curve was prepared using 2-aminoethanol (Wako, Japan), and the residual amino groups were calculated using the calibration curve. The introduction of UPy to TG was confirmed by ^1^H-NMR (400 MHz, DMSO-d6). TGUPy-42: δ = 5.73 (s, 1H, C<u>H</u>=C–C*H*_3_), 3.11 (m, 2H, NH–(C=O)–NH–C<u>H</u>_2_), 2.96 (m, 2H, C<u>H</u>_2_–NCO), 2.10 (s, 3H, C<u>H</u>_3_), 1.44 (m, 4H, NH–C*H*_2_–C<u>H</u>_2_), 1.33 (m, 4H, C*H*_2_–C*H*_2_–C<u>H</u>_2_–C<u>H</u>_2_–C*H*_2_–C*H*_2_– NCO) ppm. TGUPy-C4: δ = 5.71 (s, 1H, C<u>H</u>=C–C*H*_3_), 3.14 (m, 2H, NH–(C=O)–NH–C<u>H</u>_2_), 3.00 (m, 2H, C<u>H</u>_2_–NCO), 2.09 (s, 3H, C<u>H</u>_3_), 1.41 (m, 4H, NH–C*H*_2_–C<u>H</u>_2_) ppm. TGUPy-C8: δ = 5.73 (s, 1H, C<u>H</u>=C–C*H*_3_), 3.12 (m, 2H, NH–(C=O)–NH– C<u>H</u>_2_), 2.94 (m, 2H, C<u>H</u>_2_–NCO), 2.08 (s, 3H, C<u>H</u>_3_), 1.44 (m, 4H, NH–C*H*_2_–C<u>H</u>_2_), 1.31 (m, 8H, C*H*_2_–C*H*_2_–C<u>H</u>_2_– C<u>H</u>_2_–C<u>H</u>_2_–C<u>H</u>_2_–C*H*_2_–C*H*_2_– NCO) ppm. Fourier transform infrared (FT-IR) spectroscopy (ALPHA II, Bruker, USA) was used to check the disappearance of peak at 2280 cm^-1^ ascribing to isocyanate groups.

### 3. Rheological measurement

Rheological measurements were performed using a rheometer (MCR301, Anton Paar GmbH, Austria). SG, TG, and TGUPy-26, -42, and -53 were dissolved in phosphate buffered saline (PBS, 20 wt%) at 50 °C. The solutions were placed on the stage of the rheometer (pre-warmed to 50 °C) using a pipette, and a jig with a 10 mm diameter was set at a gap of 1 mm. After removing the excess solution, the temperature was decreased to 10 °C and started temperature sweep measurement from 10 to 50 °C at 1 °C /min. The measurements were performed at a frequency of 10 rad/s with a 1% strain in an oscillatory mode. The sol-gel transition temperature was estimated from an intersection point of the curves of storage modulus (*G’*) and loss modulus (*G”*).

To measure the shear viscosity, each solution was placed on the stage (pre-warmed to 50 °C), and a jig with a 10 mm diameter was set at a gap of 1 mm. After removing the excess solution, the temperature was decreased to 30 °C at 1 °C/min to monitor the shear viscosity. The measurements were performed at a 1% strain in a rotation mode.

### 4. Injectability test

TGUPy-42 solution was warmed up to 45 °C and loaded into a 1 mL lock-type syringe. After 1 h of incubation at 45 °C, the syringes, along with 18, 21, and 23-gauge needles, were used for measurement in a compression-testing machine (EZ-LX, Shimadzu, Japan). Each syringe was subjected to increasing stress until 50 N at 100 mm/min of approaching speed. The ejected gel was weighted, and the injectability was calculated as follows:

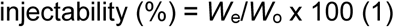

where *W*_o_ is the original weight of the adhesive before the ejection and *W*_e_ is the weight of the ejected gel.

### 5. Swelling ratio

The swelling ratio of TGUPy gel was measured by immersing in PBS at 37 °C. The swelling ratio of SG and TG were not measured because these dissolved in PBS during the swelling. TGUPy-42 were dissolved in PBS (20 wt%) at 50 °C. The solutions (100 μL) were added to a 2 mL of tube. After gelation at 25 °C for 1 h, the gels were immersed in PBS (pH = 7.4) and incubated for 24 h at 37 °C. After incubation, the gels were collected and weighed (*W*_s_). The swelled gels were then desalted by incubating in water for 24 h, freeze-dried, and weighed (*W*_d_). The swelling ratio was calculated using the following equation:

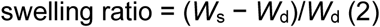

where *W*_s_ and *W*_d_ represent the weight of swelled and dried gels, respectively.

### 6. Tensile and compression test

The tensile strength of the gels was measured using a tensile tester (Shimazu) equipped in an incubator under humid conditions. SG, TG, and TGUPy-42 were dissolved in PBS (20 wt%) at 50 °C. The solutions were poured into a dumbbell-shaped silicone mold (total length: 35 mm; width: 2 mm; thickness: 1 mm; ISO 37-2). After the incubation at 25 or 37 °C for 1 h, gels were released from the mold and fixed with a 1 N clump and incubated at 25 or 37 °C. The initial distance between the clamps was set to 18 mm. Tensile tests were performed at a speed of 100 mm/min at 25 or 37 °C.

The compression test was performed using a texture analyzer (TA-XT2i, Stable Microsystems, UK). SG, TG, and TGUPy-42 were dissolved in PBS (20 wt%) at 50 °C. The solutions were poured into a silicone mold (diameter: 8mm, height: 10 mm). After the incubation at 25 or 37 °C for 1 h, gels were placed on a hot plate (pre-warmed to 37 °C) which was equipped to the texture analyzer. Gels were compressed at 100 mm/min using a jig with a 25 mm diameter to measure the stress (Maximum loading stress: 50 N).

### 7. Underwater stability test

SG, TG, and TGUPy were dissolved in PBS (20 wt%) at 50 °C. The solutions (100 μL) were added to a 2 mL of tube. After gelation at 25 °C for 1 h, PBS (1 mL) with or without 1 mg/mL of collagenase (120 U/mL, Nacalai Tesque, Inc., Japan) was added and incubated at 37 °C. After the incubation for 1, 4, 8, and 24 h, the supernatants were discarded. Ultrapure water was added to resulting gels for desalination and incubated at 25 °C for 1 h. After removing the supernatants, resulting gels were freeze-dried and weighed.

### 8. Permeability test

SG, TG, and TGUPy-42 were dissolved in PBS (20 wt%) at 50 °C. The solutions (100 μL) were added to a cell culture insert (24 well, PET, 0.4 μm pore, Corning, USA). After gelation at 25 °C for 1 h, PBS (1 mL) was added to a well plate (below inserts) and 100 μL of 100 μg/mL of fluorescein isocyanate (FITC)-labeled bovine serum albumin (FITC-BSA, Thermo Fisher Scientific, USA) was added into an insert and incubated at 37 °C. The 50 μL of the solutions were collected from lower chamber at each time point. The fluorescent intensity was measured by a microplate reader (Spark10M, TECAN, Switzerland). The dye transfer was calculated using the calibration curve.

### 9. Circular dichroism (CD) spectroscopic measurement

SG, TG, and TGUPy-42 were dissolved in PBS (0.1 wt%) at 50 °C and kept at 4 °C until the measurement. The measurement was performed using a cell with 2 mm optical path at 4 °C by CD spectrometer (J-725, JASCO, Japan). The peak at 221 nm ascribing to collagen triple helices was monitored to compare each gelatin.

### 10. Blocking test

Blocking test of hydrogen bonding was performed by evaluating the change of rheological property when competitive inhibitor added. TG and TGUPy-42 were dissolved in PBS (20 wt%, pH = 5.6) at 50 °C. By adding 1 N NaOH, the adhesive with 0.1 M NaOH was prepared (final pH was 12). Moreover, urea (Wako, Japan) was added to TGUPy adhesive to prepare the solution containing 1M urea. The adhesives were placed on the stage of the rheometer (pre-warmed to 50 °C) using a pipette, and a jig with a 10 mm diameter was set at a gap of 1 mm. After removing the excess solution, the temperature was decreased to 10 °C and started the temperature sweep from 10 to 50 °C at 1 °C /min. The measurements were performed at a frequency of 10 rad/s with a 1% strain in an oscillatory mode.

### 11. Adhesion and anti-adhesion test

Tissue adhesive properties were evaluated by measuring the adhesion strength using a Texture Analyzer (TA-XT2i, Stable Microsystems, UK) according to the American Society of Testing and Materials (ASTM) procedure (ASTM F-2258-05, Standard Test Method for Strength Properties of Tissue Adhesives in Tension). A fresh porcine-derived large intestine was used as a tissue model. The tissues were dissected into 2.5 × 2.5 cm squares and pre-warmed to 37 °C in an incubator for 1 h. The dissected tissues were bound to the probe (top) and the hot plate (bottom) with a cyanoacrylate adhesive (Loctite, Henkel, Germany). The temperature of the hot plate was kept at 37°C. Next, TG and TGUPy-42 solution (20 wt%) were warmed up to 50 °C and placed on the tissue at the bottom using a pipette. For adhesion test, immediately after adding adhesive, the top probe with tissue approached the stage at a 10 mm/minute tracking speed and pressed the tissue at 2 N (3.2 kPa) for 3 min. For anti-adhesion test, after adding adhesive, the tissue with adhesive was incubated at 37 °C for 10 min for complete gelation of adhesive. The tissue at the bottom was then pressed with top probe with tissue at 2 N for 3 min. The top probe returned to the original position at a speed of 10 mm/minute to measure the adhesion strength. After the measurement, samples were fixed in a 10% formalin neutral buffer solution and stained with hematoxylin and eosin (HE) for histological observation. The images of HE-stained tissues were scanned using a digital slide scanner (NanoZoomer S210, Hamamatsu Photonics, Japan). The temperature of adhesives on tissues were monitored using a thermography camera (FLIR ONE Pro, FLIR Systems, USA).

### 12. Cell culture

Cytotoxicity assays were carried out using a mouse fibroblast cell line (L929, RIKEN, Japan). L929 cells were cultured in RPMI1640 medium (Sigma-Aldrich, USA) supplemented with 10% fetal bovine serum (Sigma-Aldrich, USA) and 1% penicillin streptomycin (Thermo Fisher Scientific, USA). TG and TGUPy-42 were dissolved in PBS (20 wt%) at 50 °C and diluted to the range of 0.3-20 mg/mL with RPMI1640 media at 37 °C. L929 cells were seeded at 1 × 10^4^ / well in a 96-well plate and cultured for 24 h at 37 °C in a 5% CO_2_ incubator. The solutions were warmed up to 37 °C and added to each well, and cell culture continued for 24 h. The morphology of the cells was observed using an optical microscope (EVOS® XL Cell Imaging System, Thermo Fisher Scientific, USA). Cell numbers were counted using a cell counting kit (WST-8 assay, DOJINDO, Japan). Briefly, 10 μL of WST-8 reagent was added to 100 μL of culture media and incubated for 2 h. The absorbance of the media at 450 nm was monitored using a microplate reader. Cell numbers were calculated using a standard curve.

### 13. Biocompatibility and biodegradability test

All animal experiments using mice and rats were performed with the approval of the Animal Care and Use Committee of the National Institute for Materials Science. SG, TG, and TGUPy-42 were dissolved in sterile PBS (20 wt%) at 50 °C. The solutions (100 μL) were placed in a silicone mold with 1 mm thickness and 8 mm diameter and incubated at 25 °C for complete gelation in a clean bench. Gels were sterilized by UV irradiation for 30 min. Mice (7-week-old female C57BL/6J, Charles River Japan Inc.) were anesthetized by inhalation of 2% isoflurane. Hair was shaved from the backs of mice, and the shaved areas were disinfected with 70% ethanol. Gels were then subcutaneously implanted in the prepared backs of the mice. At 1, 3, and 7 days after implantation, the mice were euthanized by blood removal. For histological observation, the collected tissues were fixed in 10% formalin buffer solution for 3 days and sectioned. The images of HE-stained tissues were scanned using a digital slide scanner.

### 14. Abdominal adhesion model test

The anti-adhesion property of adhesive was evaluated using an abdominal wall-cecum defect models of rats according to previous reports [4,5]. TG and TGUPy-42 were dissolved in sterile PBS (20 wt%) at 50 °C and adhesives were sterilized by UV irradiation for 30 min at 25 °C. Adhesives were warmed up to 45 °C until the use. Rats (7-week-old male Sprague-Dawley, Charles River Japan Inc.) were anesthetized by inhalation of 2.5% isoflurane. Hair was shaved from the abdomen of rats, and the shaved areas were disinfected with 70% ethanol. The peritoneum was opened by a 5 cm long incision on the abdominal wall. The cecum was abraded using sterile surgical gauze until the tissue surface was damaged and oozing was observed, but not perforated. To damage the surface tissue of abdominal wall, a sterile silicone sheet (1 × 2 cm) was placed onto the abdominal wall and the tissue was dissected using scalpel with the fixed size. After hemostasis of damaged cecum and abdominal wall, each treatment was performed. The adhesives were injected on the damaged tissues and incubated for 5 min for gelation. For the film group, Seprafilm (hyaluronic acid/carboxymethyl cellulose film, Baxter, USA) was cut to 2 × 3 cm and was placed to damaged abdominal wall without suturing and swelled with saline. For the control group, the defects were not covered with any materials. After the injection of amikamycin (1 mg/kg), the abdominal wall was closed with 3-0 nylon sutures, and the skin was closed with a stapler. At day 14 after the treatment, rats were anesthetized by inhalation of 2.5% isoflurane, and the abdominal adhesion was checked. The severity of adhesion was classified as previously described [6]: score 0: no adhesions, score 1: mild adhesions, score 2: localized moderate adhesions, score 3: moderate and wide adhesions, score 4: severe adhesions, impossible to separate. Rats were then euthanized by blood removal, and the cecum and abdominal wall were collected. The obtained tissues were fixed in 10% formalin buffer solution for 3 days, embedded in paraffin, sectioned, and stained with HE, Masson’s trichrome, mesothelin antibody (28001, Immuno-Biological Laboratories, Japan). The images of the tissues were scanned using a digital slide scanner.

### 15. Statistical analysis

The results are expressed as the mean ± SD. One-way ANOVA, followed by Tukey’s multiple comparison post hoc test, was used to test differences among groups. The experiments were repeated multiple times as independent experiments. The data shown in each figure are a complete dataset from one representative, independent experiment. No samples were excluded from the analysis. Statistical significance is indicated as \**P* < 0.05, \*\**P* < 0.01, \*\*\**P* < 0.001, and \*\*\*\**P* < 0.0001. Statistical analyses were performed using GraphPad Prism v.8.0 (GraphPad Software).

## Acknowledgements

We appreciate the financial support from the Japan Society for the Promotion of Science (JSPS) KAKENHI (grant nos. 20K20207 and 20H02470) and the Project for Translational Research program; Strategic PRomotion for practical application of INnovative medical Technology from the Japan Agency of Medical Research and Development (AMED) (grant nos. JP20lm0203114h0001 and JP20lm0203010).

