## Supporting Information for "Hotmelt tissue adhesive with supramolecularly-controlled sol-gel transition for preventing postoperative abdominal adhesion"

**Abstract:** Postoperative adhesion is a serious and frequent complication, but there is currently no reliable anti-adhesive barrier available due to low tissue adhesiveness, undesirable chemical reactions, and poor operability. Here, we report a single-syringe hotmelt tissue adhesive to prevent postoperative abdominal adhesions. Through the augmentation of intermolecular hydrogen bonding by conjugation of the ureidopyrimidinone unit to tendon-derived gelatin, the sol-gel transition behavior of gelatin was supramolecularly-controlled, which provided a hotmelt tissue adhesive that dissolves upon warming over 40 ºC and glues at 37 ºC. This functionalization improved the key features necessary for an anti-adhesive barrier, including bulk mechanical strength, tissue adhesive properties, underwater stability, and anti-adhesive property. This hotmelt tissue adhesive with excellent tissue adhesiveness, biocompatibility, and operability has enormous potential to prevent postoperative complications.

**Table S1.** Characteristics of gelatin derived from Alaska pollock, tilapia, bovine skin, porcine skin, and porcine tendon.

| Gelatin type | Melting point of gelatin (°C) | Pro content (in 1000 amino acid residues) | Hyp content (in 1000 amino acid residues) | Pro＋Hyp (in 1000 amino acid residues) | Reference |
|---|---|---|---|---|---|
| Alaska pollock | 21.2 | 95 | 55 | 150 | (7) |
| Tilapia | 25.8 | 119 | 79 | 198 | (7) |
| Bovine, skin | 33.8 | 123 | 96 | 219 | (8,9) |
| Porcine, skin gelatin (SG) | 33.1[a] | 132 | 91 | 223 | (7) |
| Porcine, tendon gelatin (TG) | 38.2[a] | 126[b] | 92[b] | 218[b] | — |

[a] Melting points of SG and TG was measured using a rheometer. The 20 wt % gelatin solution in PBS was used. The melting points were determined from an intersection point of the curves of *G*' and *G*". [b] The contents of Pro and Hyp in TG was measured using an ion chromatography.

**Table S2.** Synthesis of TGUPy with different D.S. and spacer length of UPy unit.

| | TG | Amino groups in gelatin[a] | Spacer | UPy | D.S.[a] | Yield | Solubility in PBS |
|---|---|---|---|---|---|---|---|
| | (g) | (μmol/g) | | (mol % to amine) | (%) | (%) | |
| TGUPy-26 | 1 | 312 | C6 | 30 | 26 | 74 | ○ |
| TGUPy-42 | 1 | 312 | C6 | 45 | 42 | 72 | ○ |
| TGUPy-53 | 1 | 312 | C6 | 60 | 53 | 79 | ○ |
| TGUPy-79 | 1 | 312 | C6 | 90 | 79 | 70 | × |
| TGUPy-88 | 1 | 312 | C6 | 120 | 88 | 72 | × |
| TGUPy-C4 | 1 | 312 | C4 | 45 | 46 | 81 | ○ |
| TGUPy-C8 | 1 | 312 | C8 | 45 | 44 | 70 | ○ |

[a] The amino groups in as-prepared gelatin and degree of substitution (D.S.) was calculated by determining the residual amino groups using 2,4,6,-trinitrobenzensulfonic acid (TNBS).

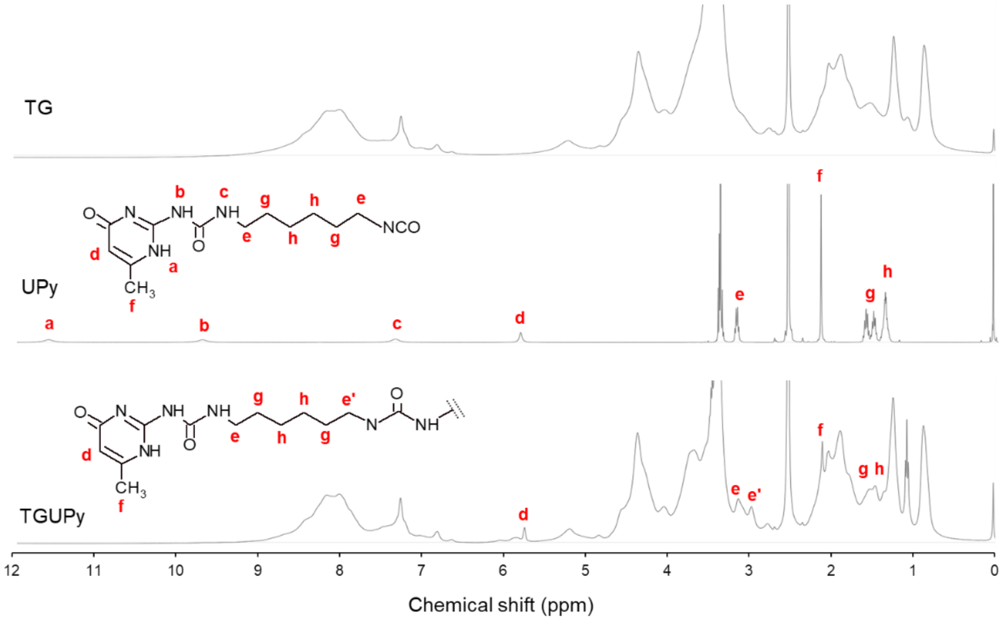


**Figure S1.** $^1$H NMR spectra of TG, UPy, and TGUPy (400 MHz, DMSO-$d_6$).

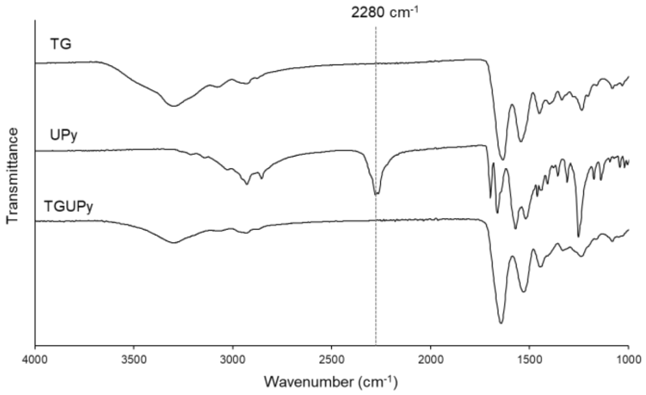


**Figure S2.** FT-IR spectra of TG, UPy, and TGUPy.

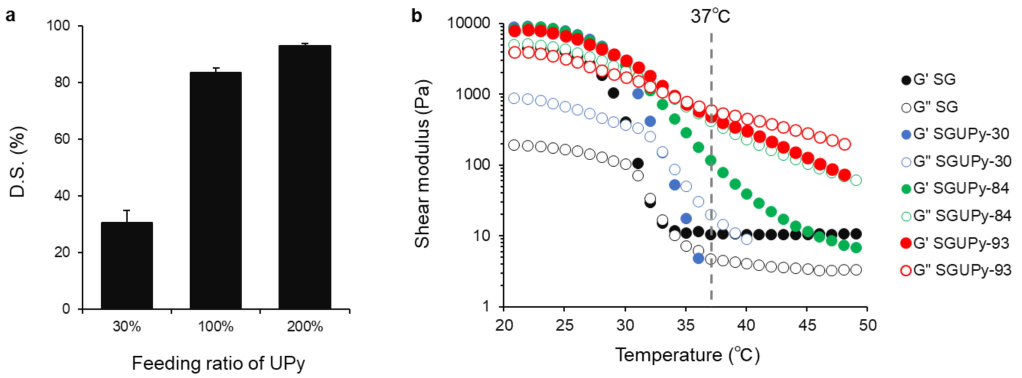


**Figure S3.** (a) D.S. of UPy in SGUPy. SGUPy was synthesized via the reaction of SG with UPy unit in the manner as TGUPy. Feeding ratio of 30, 100, and 200 UPy provided SGUPy-30, -84, and -93. (b) Temperature dependent-rheological properties of SGUPy (20 wt % in PBS) (1% strain, 10 rad/s).

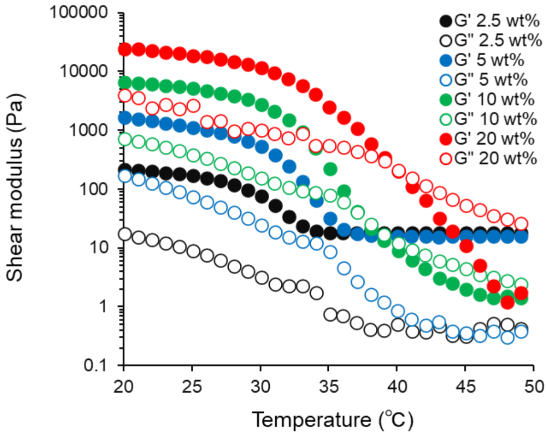


**Figure S4.** The effect of the concentration of TGUPy-42 (2.5-20 wt% in PBS) on rheological property of gels (1% strain, 10 rad/s, 37 ºC).

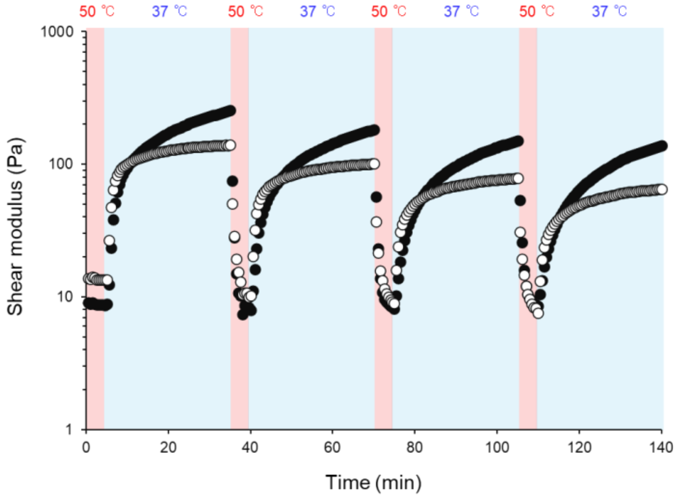


**Figure S5.** Cyclic temperature change between 50 ºC and 37 ºC in TGUPy-42 (20 wt % in PBS) in an oscillatory mode (1% strain, 10 rad/s). Each duration of measurements at 50 ºC and 37 ºC were 5 min and 30 min.

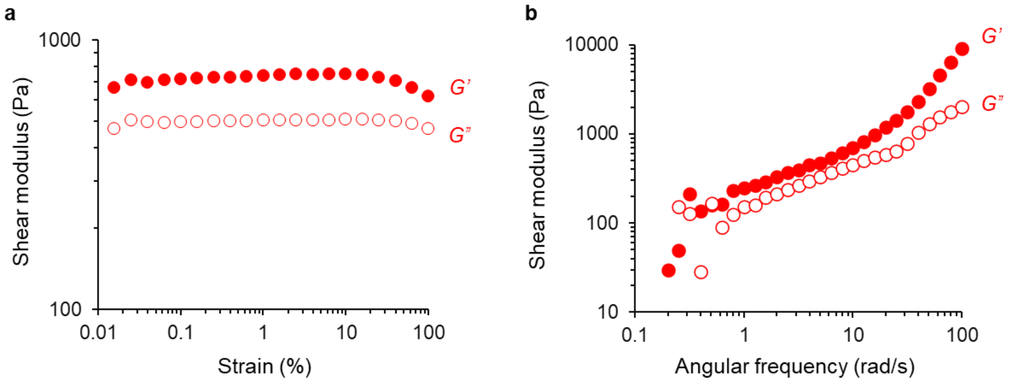


**Figure S6.** Rheological measurement of TGUPy-42 (20 wt% in PBS) as a function of (a) strain (10 rad/s angular frequency, 37 ºC) and (b) angular frequency (1% strain, 37 ºC).

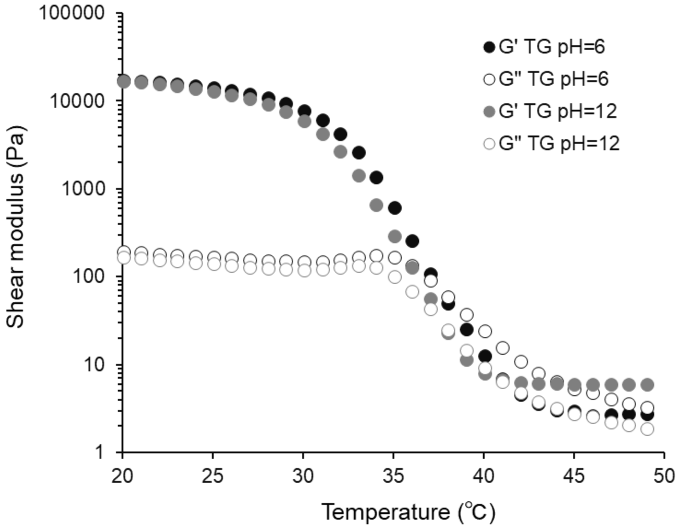


**Figure S7.** Temperature-dependent shear modulus of TG (20 wt%, pH=6, 12 in PBS) (1% strain, 10 rad/s).

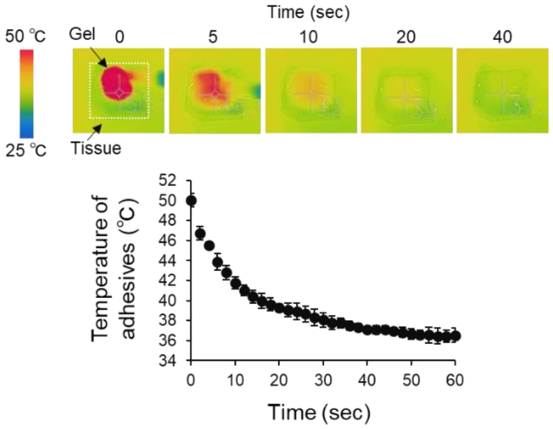


**Figure S8.** Temperature change of TGUPy-42 adhesives (20 wt% in PBS) on porcine large intestine tissue. The adhesive warmed up to 50 ºC was placed on the tissues and the temperature was monitored using a thermography camera.

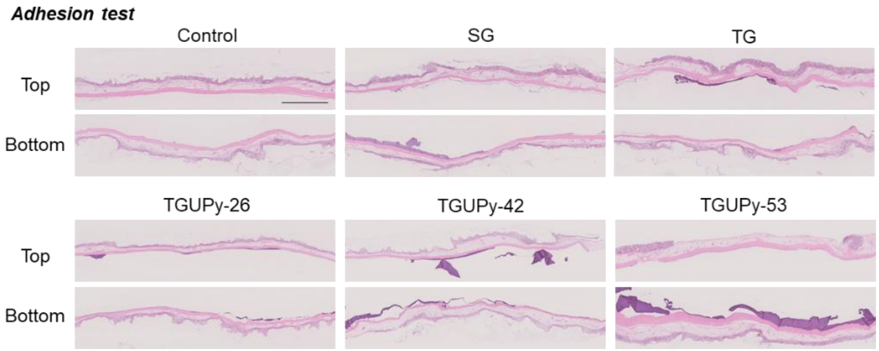


**Figure S9.** Whole images of HE-stained porcine large intestine tissues after adhesion test using SG, TG, TGUPy-26, TGUPy-42, and TGUPy-53. Control denotes the adhesion test of tissues without gels. Scale bar represents 2.5 mm.

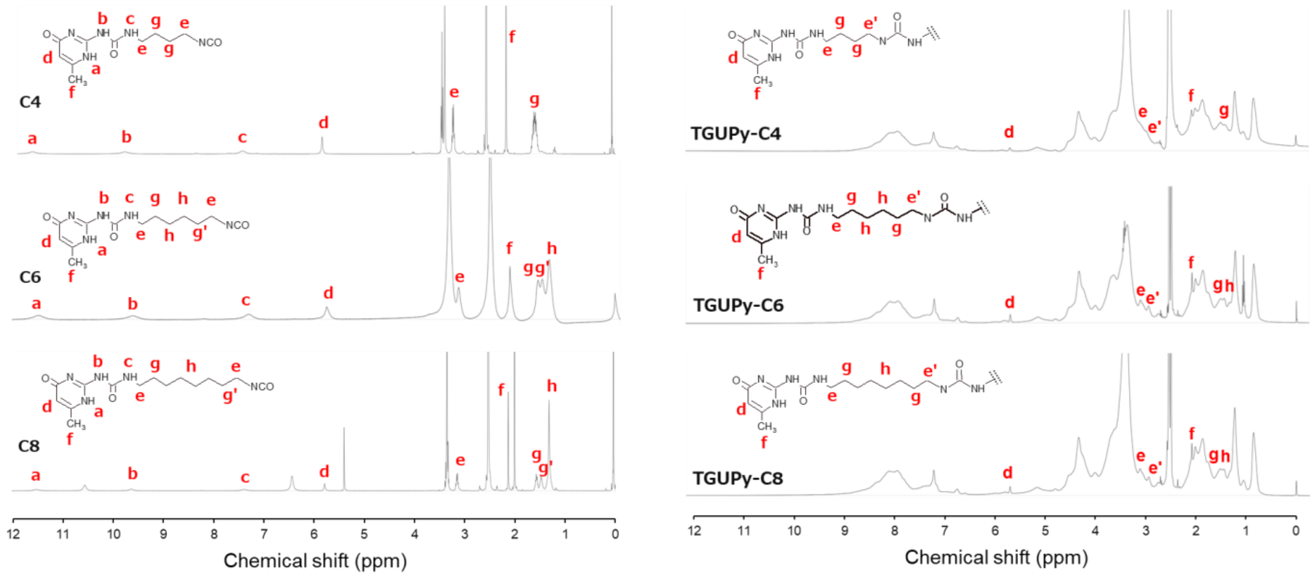


**Figure S10.** $^{1}H$ NMR spectra of C4, C6, C8, TGUPy-C4, TGUPy-C6, and TGUPy-C8 (400 MHz, DMSO-$d_6$).

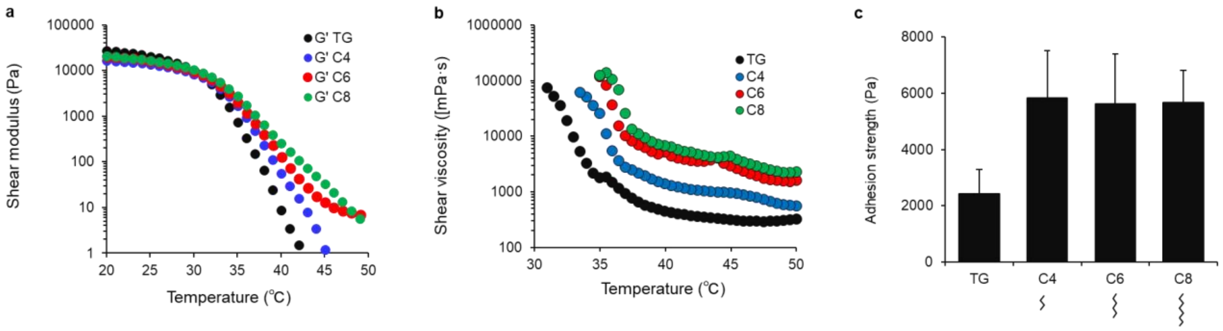


**Figure S11.** (a) Temperature-dependent shear modulus (1% strain, 10 rad/s), (b) temperature-dependent shear viscosity, and (c) adhesion strength of TG, C4-, C6-, and C8-TGUPy (20 wt% in PBS).

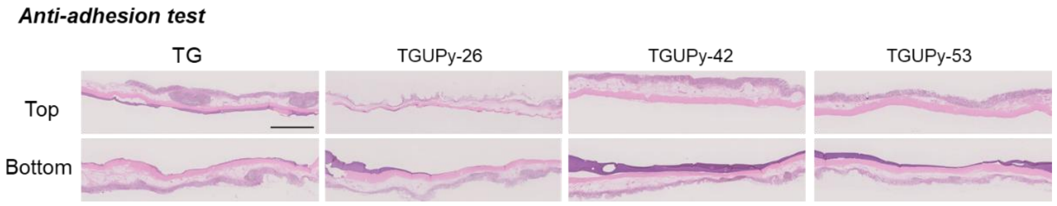


**Figure S12.** Whole images of HE-stained porcine large intestine tissues after anti-adhesion test using TG, TGUPy-26, TGUPy-42, and TGUPy-53. Scale bar represents 2.5 mm.

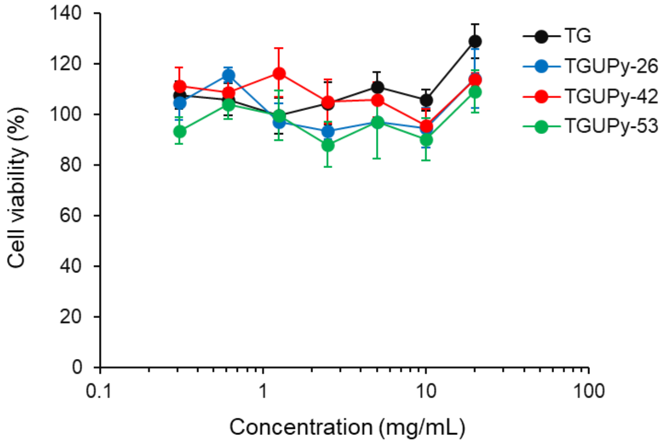


**Figure S13.** Cell viability of L929 cells exposed to media containing TG, TGUPy-26, TGUPy-42, and TGUPy-53 at 0.3 - 20 mg/mL. Cell viability was measured by WST-8 assay (n=3).

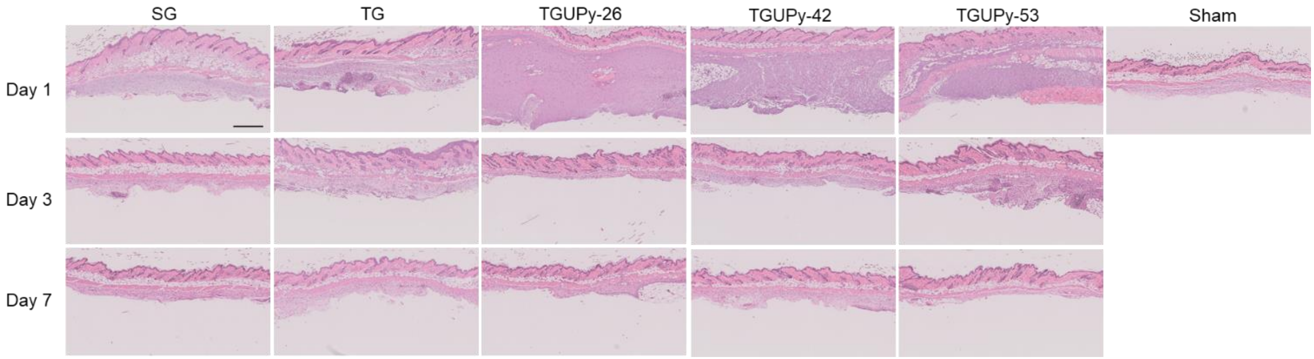


**Figure S14.** HE images of gels at day 1-7 after subcutaneous implantation in mice. The 20 wt % pre-formed gels (SG, TG, TGUPy-26, TGUPy-42, and TGUPy-53) were subcutaneously implanted in mice. TGUPy showed higher stability compared to SG and TG and all gel was degraded within 7 days.


### Author Contributions

A.N. designed and carried out the studies and wrote the paper. H.I, S.I., K.N. supported animal experiments. T.T. supervised the project and edited the paper. All authors reviewed the manuscript.
